# Shifting Routes: Plant Specific Insert trafficking and function in Arabidopsis seedlings under abiotic stress

**DOI:** 10.1101/2024.09.04.611195

**Authors:** Inês Moura, João Neves, Ana Séneca, José Pissarra, Susana Pereira, Cláudia Pereira

**Author notes:** **Correspondence:** Cláudia Pereira.

## Abstract

Due to plants’ inability to escape adverse conditions, they must adapt and adjust their endomembrane system through protein sorting and distribution. Cardosins A and B are key models for studying intracellular trafficking. They are aspartic proteinases in thistle flowers that mediate different vacuolar pathways despite sharing high sequence similarity, and both are responsive to stress conditions. The Plant Specific Insert (PSI) is a 100 amino acid domain found in these proteins. It is known that stress can impact protein sorting, shifting it from the conventional pathway (ER-Golgi) to a Golgi-independent route. In this work we assessed changes in the expression and localization of PSI from Cardosin B (PSI B) in Arabidopsis plants overexpressing PSI B-mCherry submitted to different abiotic stress conditions (saline, hydric, oxidative, metals). Aside from potential PSI B localization changes, we focused on characterizing the homozygous line, alongside assessing several biometric parameters and biochemical endpoints. The results revealed that the PSI B line responded differently depending on the stress conditions. Biometric and biochemical analyses emphasized the roles of PSI B in enhancing plant fitness and supporting adaptation to abiotic stress. Besides, confocal microscopy allowed us to find PSI B accumulation in Endoplasmic Reticulum-derived vesicles (ER bodies), indicating a shift from the common PSI B-mediated route. These findings underscore the role of PSI B in enhancing plant fitness and adaptation to abiotic stress through altered protein trafficking.

**Highlight:** PSI B has an active role in enhancing plant fitness, revealing its value in adaptation and tolerance to abiotic stress by adjusting its localization and trafficking under challenging environments.

## 1. Introduction

Plants’ natural environment consists of a complex interaction of abiotic and biotic factors, and as sessile beings, plants face the critical challenge of coping with, and adjusting to, the environmental conditions. Those two factors frequently emerge as key stresses for plants, with the capacity to act separately or, more usually, in combination [1,2]. Drought, salinity, and high temperatures are examples of such factors [3,4]. We are currently living in an era marked by notable climate change, which presents a considerable challenge to plant development and agricultural yield. Consequently, it becomes crucial to investigate the fundamental biological aspects associated with how plants perceive stress signals and respond to challenging climatic scenarios. Plants have evolved the ability to acclimate to dynamic conditions such as physiological adjustments and molecular responses. These mechanisms include changes in transcription and translation, as well as post-translational modifications, the regulation of protein degradation and accumulation [5–7]. The plant endomembrane system is a highly sophisticated and organized network that includes the transport of molecules through vesicles via secretory and endocytic routes. Notably, this fundamental mechanism allows adjustments and adaptations during stress, ensuring the maintenance of homeostasis [8]. One of the most activated processes during stress response is autophagy that plays a critical role in increasing tolerance to oxidative stress, which is typically linked with a range of stress conditions [9]. This process is induced in response to various stresses, including osmotic and saline stresses [10,11]. Other mechanism of plant’s response to stress, are the formation of ER bodies (ER-derived compartments) that contribute as another vital process in the endomembrane response. They are involved in the unconventional trafficking to the vacuole and are typically inactive in healthy cells, becoming actively involved in cell death, when cells are stressed or injured [12]. Hayashi and collaborators reported that during stress the number of ER bodies that direct fuse with the vacuole increases [13]. Stress can disrupt the conventional sorting of proteins into the vacuole, and cells must promptly adjust their cargo trafficking mechanisms. This adaptive response often involves following unconventional routes to efficiently meet the demands imposed by stress [14,15]. Cardosins serve as a great model for studying protein sorting throughout the endomembrane system. These are aspartic proteinases (APs), isolated from the pistils of the *Cynara cardunculus* plant, and constitute over 70 % of the entire protein content in these structures [16]. Some plant APs, have an independent domain consisting of about 100 amino acids, the plant-specific insert (PSI) [17]. The PSI’s function is not entirely understood. In 2002, Egas and colleagues [18] proved the PSI ability to interact with membranes and induce vesicle leakage. These findings suggest that this domain can cause disruption on cell membrane permeability, perhaps acting as a defensive mechanism against pathogens and contributing to cell death processes [18–21]. When isolated and *in vitro*, StAP-PSI (PSI domain of potato aspartic protease) has been shown to display antimicrobial activity against both plant and human pathogens [20,22,23] as well as anticancer activity against leukemia cells without causing toxicity [24]. Furthermore, PSI have been shown to be involved in vacuolar targeting of APs in diverse plant species [25–27]. Pereira and co-workers (2013) showed that the PSI domains present in cardosins A and B have the ability to direct secreted proteins to the vacuole through different pathways [28]. In contrast to other plant APs like Cardosin B, the PSI domain of Cardosin A lacks a glycosylation site [29]. The presence or absence of a glycosylation site on the PSI domain emerges as a significant structural element that determines the exact path that PSI-driven targeting takes [28]. As a result, when PSI is glycosylated, its transport to the vacuole becomes dependent on COPII vesicles, leading it to take the conventional route. If the PSI is not glycosylated, transport is no longer mediated through COPII vesicles, and thereby the Golgi is bypassed. In this context, this work aims to characterize an Arabidopsis line overexpressing PSI B-mCherry and understand potential modifications in PSI B subcellular localization induced by abiotic stress. Our investigation was extended to evaluate the potential plant tolerance enhancement through the overexpression of this domain under stress conditions. This objective was particularly motivated by prior research, which demonstrated that the expression of plant-specific insert of potato aspartic proteases (StAP-PSI) confers enhanced resistance to biotic stress in Arabidopsis plants [23]. Our results revealed significant differences in the response of the PSI B line under different stress conditions. Particularly noteworthy was the identification of PSI B accumulation in ER bodies, an unquestionably novel and significant discovery, particularly because ER-bodies fused directly to the vacuole, without contribution of the Golgi. This finding has triggered a whole bunch of hypotheses and uncertainties regarding PSI B’s potential to mediate a Golgi-independent pathway to the vacuole.

## 2. Materials and Methods

### 2.1 Biological Material and Stress Assays

Homozygous PSI B-mCherry lines were obtained, from *Arabidopsis thaliana (col0),* using the floral-dip method [30] and gDNA extraction was performed according to the manufacturer’s instructions using the “GRS Genomic DNA kit Plant” (Grisp) extraction kit. The presence of the transgene was performed by PCR using *mCherry* primers listed in Table 1, and Phusion High-Fidelity PCR Master Mix with HF Buffer (Thermo Scientific™). All PCR products were checked by separating in a 1 % (w/v) agarose gel in 1x TAE [40 mM Tris-HCl and 1 mM EDTA pH 8.3] with GreenSafe Premium (NZYTech). For the detection of PSI B fused with mCherry, a Western Blot was performed with a Trans-blot turbo transfer system (BIO-RAD) in accordance with the manufacturer’s instructions. The gel was transferred to a nitrocellulose membrane at 100V for 10 minutes and the membrane was then transferred to the blocking solution [5 % (w/v) skim milk and 0.5 % (v/v) Tween-20 in TBS-T [20 mM Tris, 150 mM NaCl, 0.1% (w/v) Tween®20] for 30 minutes at RT. The primary antibody, rabbit polyclonal anti-mCherry (Merk Millipore) was diluted (1:1000) in blocking solution before being incubated at 4 °C ON with steady shaking. Following that, three washings in TBS-T were conducted and 1h incubation with the secondary antibody - Goat anti-Rabbit IgG (HRP) (Merk Millipore) diluted in TBS-T (1:5000) - was performed. The membranes were washed, and signal was developed using the Clarity ECL Western Blotting Substrate Kit (BioRad), as directed by the manufacturer. The imaging was done through digital visualization with the ChemiDoc XRS+ System (BioRad).

**Table 1.**
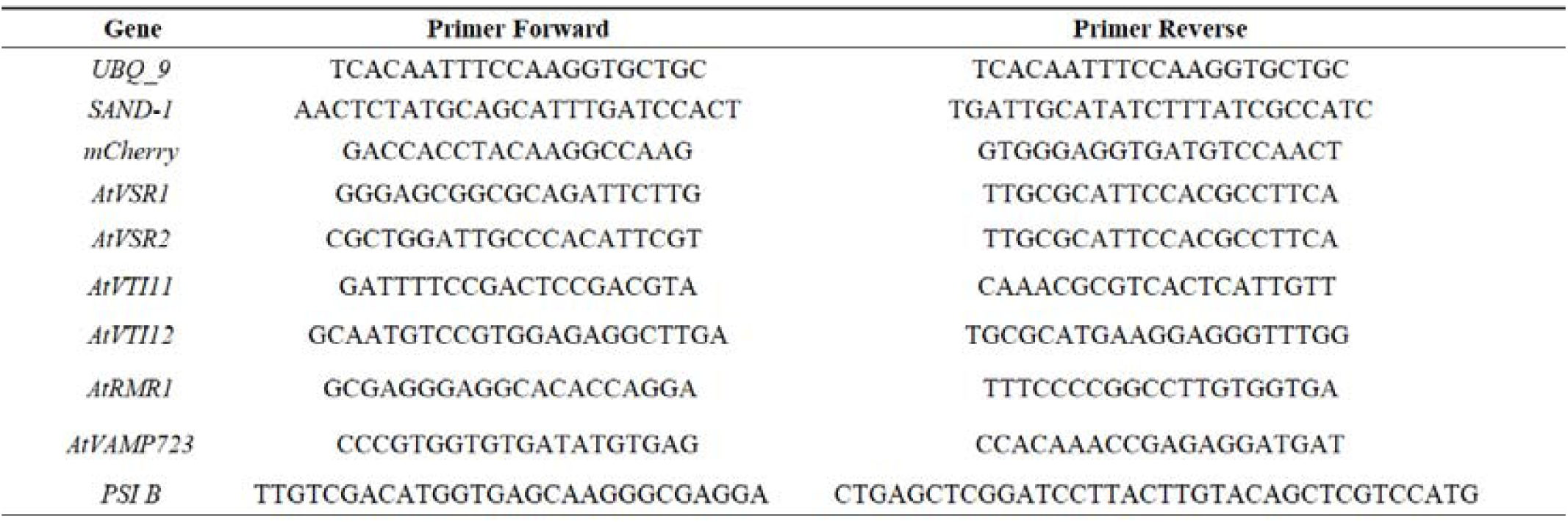
List of genes and corresponding primer pairs used in the quantitative RT-PCR assay.

Seed germination was performed in MS (Murashige & Skoog, Duchefa Biochemie) medium with 1.5 % (w/v) sucrose, 0.7 % (w/v) bactoagar, and pH 5.7. To simulate the abiotic stress conditions, different stress-inducing agents were added to the medium: sodium chloride (saline stress – 50 mM), mannitol (hydric stress - 50mM and 100mM), hydrogen peroxide (oxidative stress - 0.5mM), and zinc sulfate (metal-induced stress - 150µM). Seeds were stored at 4 °C for 2-3 days to induce stratification, and then were placed in a growth chamber for approximately 10-12 days under the following conditions: 22 °C with 60 % humidity and a 16 h light photoperiod (OSRAM L 36W/77 and OSRAM L 36W/840) at an intensity of 110 μmol/m^-2^ s. Three biological replicates were prepared.

### 2.2 Growth analysis

To assess potential phenotypic impacts of PSI B overexpression, a developmental assay was conducted based on development processes [31] comparing WT to PSI B transformed plants. Seeds were germinated on a solid medium, as previously described, and then transferred into soil during the seedling stage. The time of appearance of different developmental points was recorded for further analysis: seed inhibition; cotyledons fully opened; two rosette leaves; three rosette leaves; four rosette leaves; first flower buds visible; first flower open and presence of first siliques (Figure 1-D).

**Figure 1 –.**
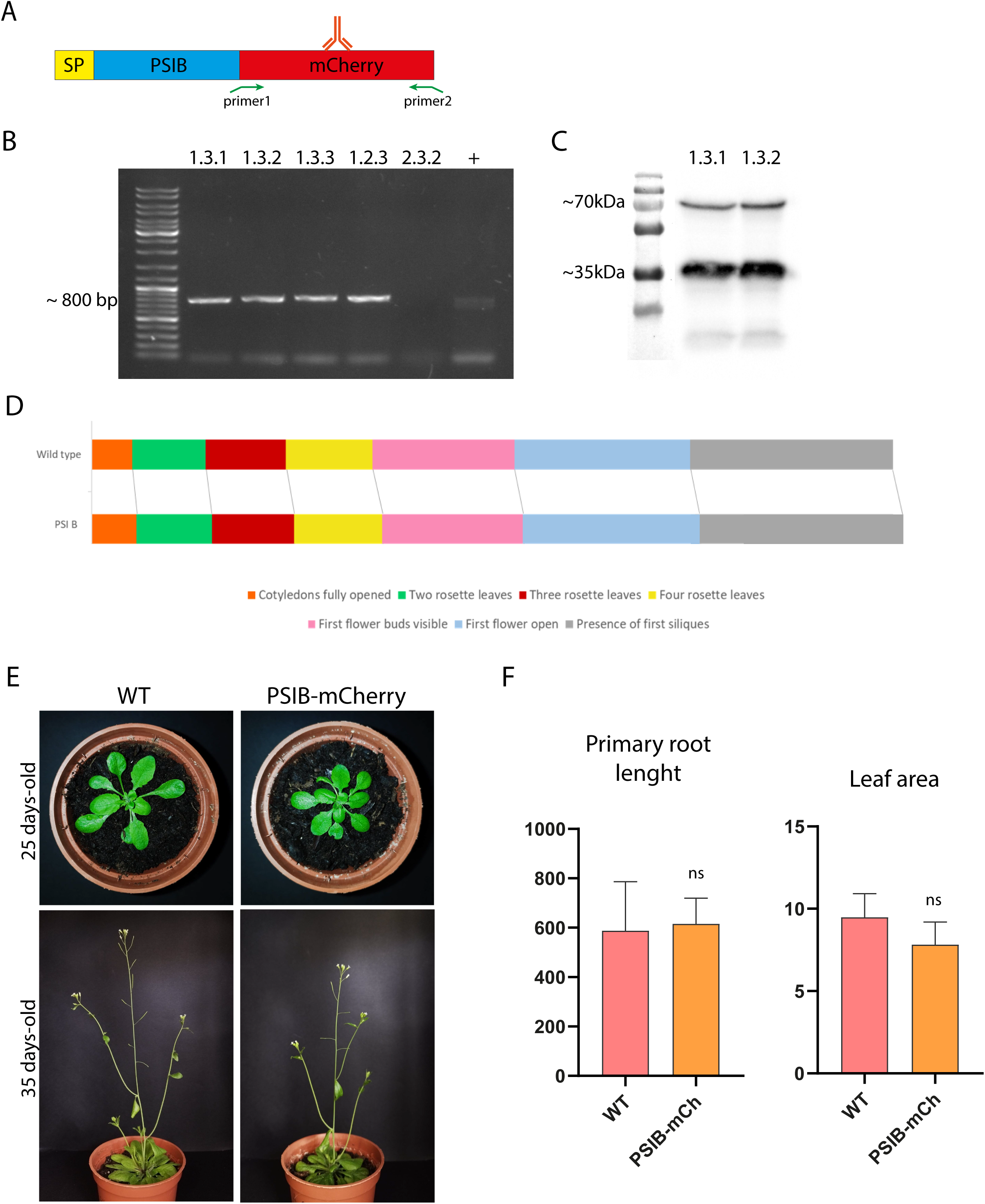
Characterization of the Arabidopsis line overexpressing SP-PSI B-mCherry. **A** - Schematic representation of SP-PSIB-mCherry construct. Y-shaped structure (upside down) representing the primary antibody used (rabbit polyclonal anti-mCherry); SP-signal peptide; **B** - Electrophoretic analysis of different gDNA’s from five T2 PSI B-mCherry lines chosen for transformation validation. Ladder: GeneRuler DNA Ladder Mix - 0.5 g/L (Thermo Fisher Scientific); + - positive control (pVK-SP: mCherry); **C** - Western blot analysis of A. thaliana protein extracts from two T2 PSI B-mCherry overexpressing lines using anti-mCherry antibody. Ladder: PageRuler™ Plus Prestained Protein Ladder (Thermo Scientific); **D** - Scheme representing the chronological progression of the 7 principal growth stages selected for phenotypical comparison between PSI B Arabidopsis line and wild-type plants; **E** - Arabidopsis thaliana plants in two developmental stages. Wt and PSI B-mCherry plants 25- and 35-days post-germination; **F** - Bar plot of the primary root length and leaf area of Wt and PSI B-mCherry plants. Results are presented as mean ± standard deviation (SD) and result from the evaluation of at least three experimental replicates (n ≥ 3). The statistically experimental from One-way ANOVA values are represented by “ns” (not significant), according with the One-way ANOVA.

### 2.3 Biometrical Analysis

A phenotypic study was carried out to determine if overexpression of PSI B may affect certain biometric parameters. After the seeds were sown the plates were stored in the conditions already described, but square plates were used and incubated vertically, to promote vertical root growth. Following 12 days post-germination, the root length and the leaf area were measured. Leave pictures were captured using a magnifying glass, and root and leaf area analysis were performed using the ImageJ/Fiji software under two conditions: Arabidopsis WT plants and selected Arabidopsis lines overexpressing PSI B-mCherry.

### 2.4 Pigment Quantification

Following the 12 days after germination, four replicates of each stress condition were collected for biochemical analysis. Seedlings (80 mg) were homogenized at room temperature with 80% (v/v) acetone. After centrifugation at 1400 x *g* for 10 minutes, the supernatant (SN) was collected and kept in the dark. Chlorophyll and carotenoid levels were determined spectrophotometrically by recording the absorbance at 470, 647, and 663 nm and calculated using the formulas outlined by Lichtenthaler (1987) [32].

### 2.5 cDNA Preparation

100 mg of seedling tissue was weighed to obtain total RNA preparations, and the “NZY Total RNA Isolation Kit” (NZYTech) was used following the manufacturer’s instructions. Total RNA was quantified using a microdrop spectrophotometer (DeNovix DS-11, Bonsai Lab), and its integrity was confirmed on a 1% agarose gel. The cDNA synthesis from RNA samples was conducted using the “NZY First Strand cDNA Synthesis Kit” (NZYTech) according to the protocol provided. Both RNA and cDNA were then stored at −80 °C.

### 2.6 Quantitative RT-PCR

The quantitative RT-PCR was performed in a CFX96 Real-Time System (Biorad) using PowerUp SYBR Green Master Mix (Thermo Fisher, Waltham, MA, USA). Three biological replicates and three technical replicates were performed for each gene and condition. In short, the 10 µL reaction comprised 400 nM of each primer (Table 1) and 2 µL of cDNA in a 1:8 dilution. The PCR reaction was conducted with the following conditions: initial denaturation (95 °C for 3 min), followed by 40 cycles of 95 °C for 10 s, 56 °C for 10 s, and 72 °C for 30 s. After 40 cycles, a melt-curve was generated employing the following conditions: 95 °C for 10 seconds, followed by a continuous temperature increase from 65°C to 95°C. The housekeeping genes SAND-1 and UBC9 were used as reference genes and all primers used have been validated in a previous study from the lab [33]. The results analysis was made by relative quantification using Bio-Rad CFX Maestro (version 1.0) software and compared with the control condition.

### 2.7 Confocal Microscopy

Arabidopsis seedlings expressing PSI B-mCherry were grown under the previously mentioned stress conditions and imaged using Confocal Laser Scanning Microscopy (CLSM, Leica SP5). The biological material (cotyledons or small sections of rosette leaves) was placed on a slide with a drop of sterile water covered by a cover slip. The lower epidermis was imaged. mCherry emission was detected between 580-630 nm, using 561 nm excitation. The ImageJ® /Fiji program was used to analyze and processed the obtained images.

### 2.7 Drug Treatment Assay

Brefeldin A (20 µg/mL), Cytochalasin (20 µM) and Oryzalin (5 µg/mL) solutions were prepare in MS liquid medium, and 2 mL of each drug was poured independently in each well of the 6-well plates, over the seedlings. Seedlings were kept in the drug treatment until the cells were imaged (4h).

### 2.8 Transmission Electron Microscopy

Small sections measuring 1-2 mm from cotyledons (12 days old) grown in different abiotic stress conditions were cut for transmission electron microscopy. For this assay both WT and PSI B-mCherry lines were fixed in Na-PIPES 1.25 % (w/v) pH 7.2 and Glutaraldehyde 2.5 % (v/v) for 1 hour at room temperature (RT). The samples were then washed with 1.25 % (w/v) Na-PIPES (3 times for 10 min each) and post-fixed in 4 % (w/v) osmium tetroxide (OsO4), prepared in 1.25 % (w/v) Na-PIPES, at RT for 1 h. After that, 3 washes with Na-PIPES 1.25 % (w/v) pH 7.2 for 10 minutes were performed. Then, the samples were subjected to a dehydration step in a graded series of increasing ethanol concentrations: 10 % (v/v) for 10 minutes; 20 % (v/v), 30 % (v/v) and 50 % (v/v) for 15 minutes; and 70 % (v/v), 90 % (v/v) and 100 % (v/v) for 20 minutes. The ethanol was then removed, and the biological material was incubated in propylene oxide for 20 minutes. The biological material was incubated in 10% resin and 90% propylene oxide for 15 minutes and drops of resin were incrementally added until the resin: propylene oxide ratio reached 50:50. The next day the resin and propylene oxide solution was removed and replaced with pure resin (overnight). The samples were then transferred to oriented molds covered with Epoxy resin and polymerized at 60 °C overnight. Ultrathin sections (60–80 nm) were acquired using a diamond knife (Diatome) in an Ultracut UC6 (Leica) ultramicrotome and placed in 400 mesh copper grids. Post-staining was performed with Uranyless EM Stain (Uranyless) and Reynolds Lead Citrate 3 % (Uranyless) for 5 min each. The grinds were then rinsed several times to remove excess stain and dried on filter paper before being examined using a Transmission Electron Microscope (TEM - Jeol JEM 1400). Images were captured with an Orius® SC200D camera and image analysis and processing were done using ImageJ®/Fiji software.

## 3. Results

### 3.1 Characterization of PSI B-overexpressing Arabidopsis line

Arabidopsis lines carrying SP-PSI B-mCherry (hereafter termed PSI B – Figure 1-A) were germinated until the T4 generation to ensure homozygosity. Correct transgene insertion was validated by PCR over genomic DNA and five independent lines were tested. The transgene – band of approximately 800 bp (corresponding to mCherry) – was observed in all lines, except for line 2.3.2 (Figure 1-B). Two different lines were selected to check the expression of PSI B fused to mCherry by Western blot and using an anti-mCherry antibody. The expression was confirmed by the presence of a band of 35 kDa, corresponding to the PSI B-mCherry protein and a fainter band of about 70 kDa, consistent with the presence of a dimer (Figure 1-C). To explore potential phenotypic implications or developmental delays induced by protein overexpression a thorough test comparing PSI B-expressing plants with WT plants was performed. The assay intended to confirm the regular succession of essential growth stages throughout Arabidopsis development. Following a 48-hour period of seed stratification, all plants were grown under the same conditions. Seven developmental stages were evaluated: cotyledons fully opened; two rosette leaves; three rosette leaves; four rosette leaves; first flower buds visible; first flower open and presence of first siliques. According to the data presented in Figure 1-D, the growth patterns of the PSI B line closely resembled those of the WT, with a slight delay in seed germination (Figure 1D-E). Moreover, quantification of primary root length and leaf area between WT and PSI B-expressing plants do not show significative differences (Figure 1F).

### 3.2 PSI B localizes to the vacuole and ER-derived bodies

To characterize and gain in-depth knowledge of the expression and localization of this protein, PSI B Arabidopsis lines were observed under a Confocal Laser Scanning Microscopy (CLSM), and different plant tissues were considered: cotyledons and rosette leaves. Figure 2-A (a and c) illustrates the expected predominant accumulation of PSI B-mCherry in the vacuole. Nonetheless, in both cotyledon and rosette leaves, it was also detected in association with the ER, as indicated by the blue arrows (Figure 2-A, b and d), sometimes visible as small aggregates. Particularly evident in cotyledons, PSI B-mCherry was detected in elongated ER-derived aggregates (Figure 2-A, d), previously identified as ER-bodies [34]. To have a more dynamic insight on the nature of these ER-bodies, BFA (blocking the ER-to-Golgi transport), Cytochalasin D (CytD – affecting the actin cytoskeleton) and Oryzalin (Ory – promoter of microtubule disruption) treatments were performed (Figure 2-B). In all cases, PSI B-mCherry was still detected in the vacuole and in the ER-derived elongated bodies. However, when applying BFA, some larger aggregates were visible (Figure 2-B, yellow arrows), indicating some effect of the drug. When applying CytD, the elongated structures could be observed clustered inside the cells (Figure 2-B, green arrows), which was not observed in the Ory treated cells. Time-lapse experiments (Figure 2C) showed that the PSI-B labelled ER-bodies moved along the ER network (Figure 2C, a – blue circles; Sup movie 1) and that movement stopped when CytD was applied (Figure 2C, b – blue circles; Sup movie 2), but still moved upon Ory treatment (Figure 2C, c – blue circles; Sup movie 3)

**Figure 2 -.**
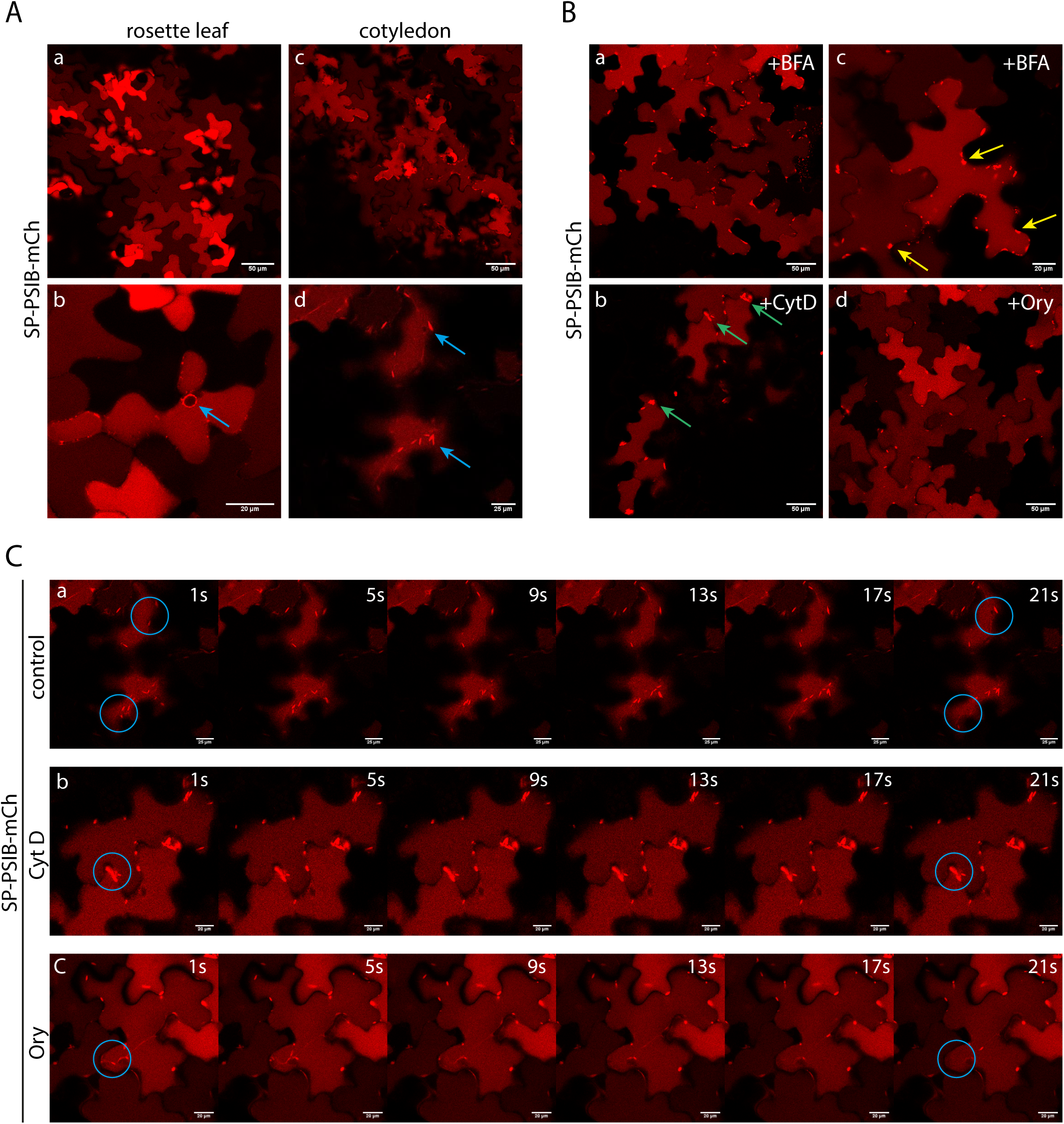
Subcellular localization of PSI B-mCherry in overexpressing A. thaliana plants. **A** - Images of PSI B-mCherry in rosette leaves (a, b) and cotyledons (c, d). In rosette leaf, the blue arrow indicates ER accumulation, and in cotyledon indicate ER bodies accumulation of PSI B-mCherry. **B** - Subcellular localization of PSI B-mCherry in cotyledons of A. thaliana line treated with BFA (a, b), Cytochalasin (c) and Oryzalin (d). Yellow arrows indicate BFA agglomerates, and green arrows indicate clustering of ER-bodies. **C** - Stills retrieved from time-lapse movie illustrating the movement of PSI B-mCherry-labelled compartment (blue circle) under control (a), Cytochalasin D (b), and Oryzalin (c) treatments. Images were acquired using the 561 nm laser for mCherry and were analysed and processed using ImageJ/Fiji software.

### 3.3 PSI B overexpressing seedlings respond to stress conditions

Since there is evidence that aspartic proteinases are associated with stress responses, abiotic stress assays were conducted to investigate the potential impact of PSI B overexpression in *A. thaliana* seedlings when subjected to adverse conditions. In a first approach, seedling morphology, average root length, and leaf area were measured in plants under stress. Concerning the seedlings’ morphological characteristics, when comparing control to stress conditions, PSI B-mCherry line displayed considerable modifications. Figure 3-A illustrates a delay in seedling development in the saline stress condition (S1), along with the two hydric stress conditions (H1 and H2). Notably, when comparing the S1 condition to H1 and H2, distinct morphological differences became evident, being H2 the harshest one, both in WT and PSI B lines (Figure 3-A). Conversely, under oxidative conditions (Ox), morphological changes were not significant, as these seedlings closely resemble the control. In contrast, the metal-induced stress by Zn led to the development of chlorotic areas on the leaves, not observed in WT seedlings. Analysis of the primary root length of Arabidopsis WT plants and PSI B-mCherry line was performed in the same stress conditions. The results regarding PSI B-mCherry line root length showed significative difference between the different stress conditions (Figure 3-C). An accentuated decrease could be found in S1, H2 and Ox conditions (Figure 3-C), similarly to what was observed (despite with milder effects) on WT plants (Figure 3B). In H1, both PSI B and W showed an increase in root length (Figure 3-B, C). When directly comparing PSI B with WT plants, it was clear an increase in primary root length of PSI B plants both in control (C) and H1 condition (Sup Figure 2-A). During the stress trials, it became evident that morphological changes in seedling development had a major influence on leaf area. As anticipated, in the case of PSI B, the conditions that led to a more pronounced delay in development also correlated with a significant reduction in leaf area – S1, H1, and H2 (Figure 3-E). The same was observed for WT (Figure 3-D). Interestingly, and contrary to what was observed for WT plants, PSI B plants presented a higher leaf area in the Ox condition (Figure 3-E). When comparing the two lines, WT plants had significantly greater leaf area than the overexpressing line, in the S1 and H2 conditions (Sup Figure 2B). In contrast, in the Ox and Zn conditions, the PSI B line reported higher values compared to WT (Sup Figure 2B).

**Figure 3 -.**
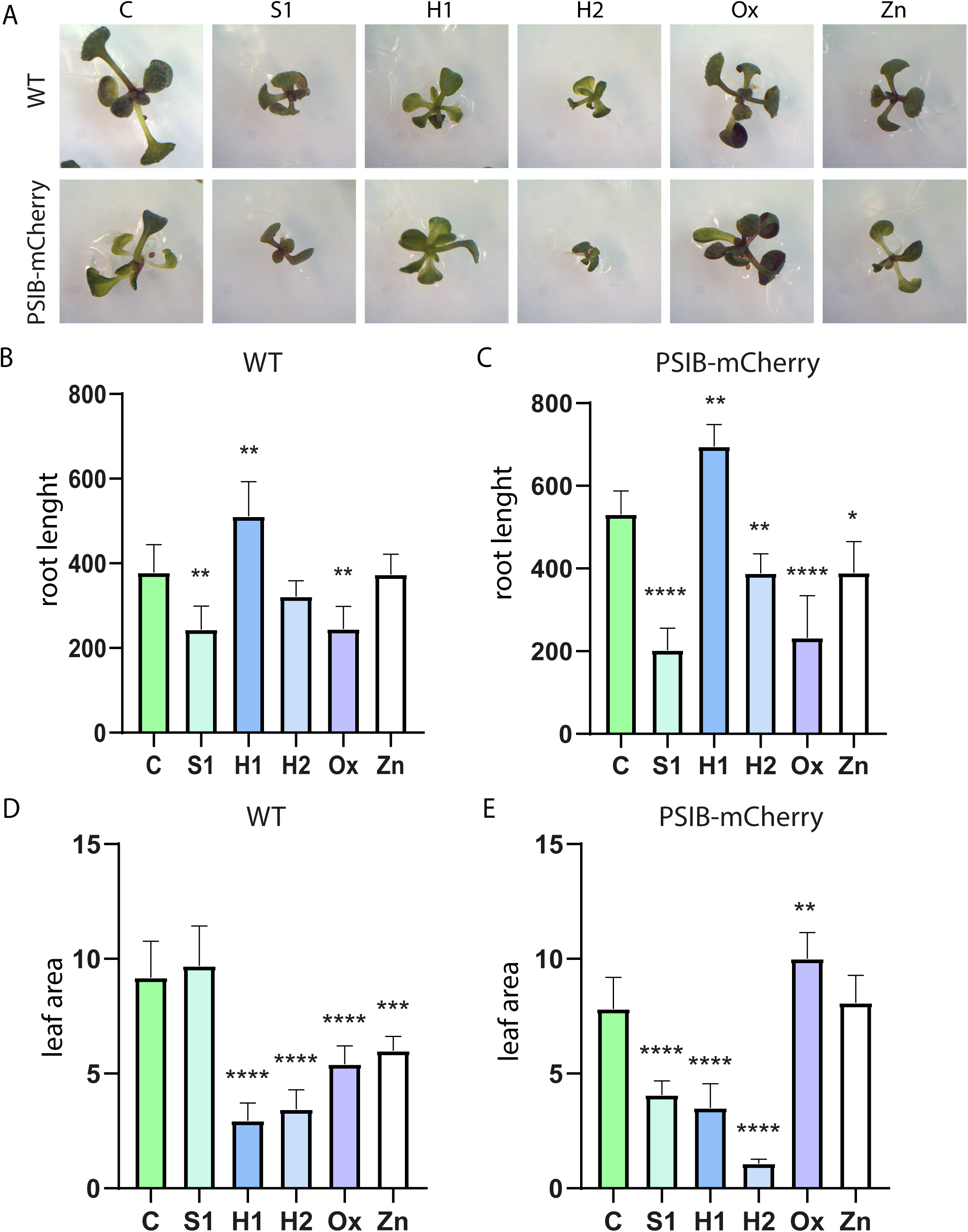
Biometric comparison of both Arabidopsis thaliana WT and PSI B-mCherry line under abiotic stress. **A** - Images of seedlings germinated for 12 days under stress conditions. (C) Control; (S1), saline stress, (H1, mild hydric stress; (H2), high hydric stress; (Ox), oxidative stress and (Zn) stress induced by metals. **B** - Bar plot of the root length measurements under stress conditions in WT seedlings; **C** - Bar plot of the root length measurements under stress conditions in PSI B lines; **D** - Bar plot representing the leaf area in WT seedlings under stress conditions; **E** - Bar plot representing the leaf area in PSI B lines under stress conditions. Results are presented as mean ± standard deviation (SD) and result from the evaluation of at least three experimental replicates (n ≥ 3). The statistically experimental from One-way ANOVA values are represented by: * (pvalue ≤0.05), ** (pvalue ≤0.01), *** (pvalue ≤0.001); **** (pvalue <0.0001), according with the One-way ANOVA.

To investigate whether the phenotypic changes previously observed, particularly at the leaf level, were reflected in biochemical parameters, we assessed the physiological performance of both WT and PSI B seedlings under abiotic stress conditions. In this context, we focused our attention on evaluating photosynthetic pigments, as they are considered relevant indicators for this study. Figures 4-A and B illustrate carotenoids content both in PSI B and WT seedlings under different stress conditions. A significant increase in carotenoids levels was observed for the H1 and H2 conditions in PSI B plants (Figure 4-B), while for WT only the H2 conditions show a significant increase (Figure 4-A). No significant alterations were found for the rest of the conditions under study. Despite the changes observed when the different stress conditions are compared with control, no significative changes were detected when comparing WT and PSI B lines, under the same condition (Sup Figure 2C). Analyzing total chlorophylls, no significant changes were observed between the different stress conditions applied to the PSI B line (Figure 4-D), while a decrease in chlorophyll content was observed in WT plants, for S1, Ox and Zn conditions (Figure 4-C). However, when directly comparing the chlorophyll content of WT and PSI B plants, and despite the increasing tendency in S1, Ox and Zn, no significative differences were recorded (Sup Figure 2-D).

**Figure 4 -.**
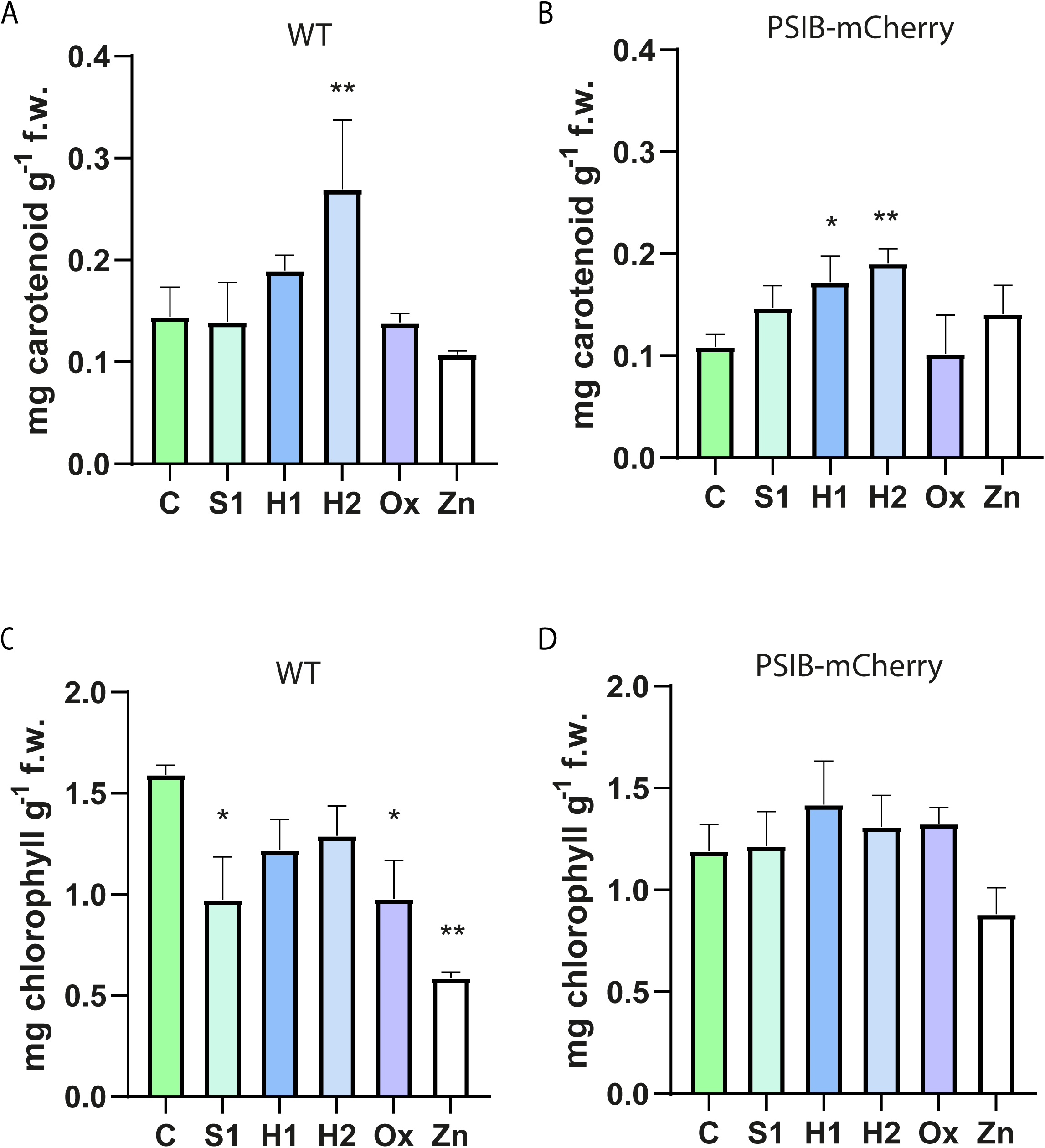
Pigment Quantification in Arabidopsis thaliana WT and PSI B-mCherry lines (seedlings germinated for 12 days) under abiotic stress. **A** - Bar plot of carotenoid quantification (mg g-1 fw) in stress conditions relative to control from WT plants. **B** - Bar plot of carotenoid quantification (mg g-1 fw) in stress conditions relative to control from PSI plants. **C** - Bar plot of the total chlorophylls (mg g-1 fw) in stress conditions relative to control from WT plants. **D** - Bar plot of the total chlorophylls (mg g-1 fw) in stress conditions relative to control from PSI plants. Results are presented as mean ± standard deviation (SD) and result from the evaluation of at least three experimental replicates (n ≥ 3). The statistically experimental from One-way ANOVA values are represented by: * (pvalue ≤0.05), ** (pvalue ≤0.01), according with the One-way ANOVA.

### 3.4. Ultrastructural analysis reveals subtle changes in endomembrane organization under stress

After checking the phenotypical parameters, our focus turned to the ultrastructural characterization of the PSI-B plants under control and abiotic stress conditions, particularly concerning subtle alterations in organelle morphology. The WT line (Figure 5-A a,b) was also imaged as it represents the normal cellular arrangement, and PSI B-mCherry cells (Figure 5-A c,d), in control condition, were compared with both the WT line and abiotic stress situations. The analysis aims at clarifying not only the differences between WT and PSI B-mCherry line, but also the impact of abiotic stress on the cell ultrastructure. The WT line demonstrated a characteristic and accurate organization of cellular organelles. It was possible to observe well-structured chloroplasts with intact thylakoid arrangements in grana stacks as well as a high number of starch grains (Figure 5-A a). The membranes were well defined, including both the cytoplasmic membrane and the membranes of different intracellular compartments. Mitochondria, Golgi and ER were all observable, and several vesicles were visible in both images (Figure 5-A a-d). Looking at cells from the control PSI B-mCherry line, it was immediately evident that the prominent feature was the abundance of starch grains, which were not only numerous but also notably larger in size (Figure 5-A c). Alongside this observation, the organization of chloroplasts was once again recognizable, although the definition of its internal membranes might not be as distinct as in WT. Nevertheless, the cellular organization is quite similar to WT cells.

**Figure 5 –.**
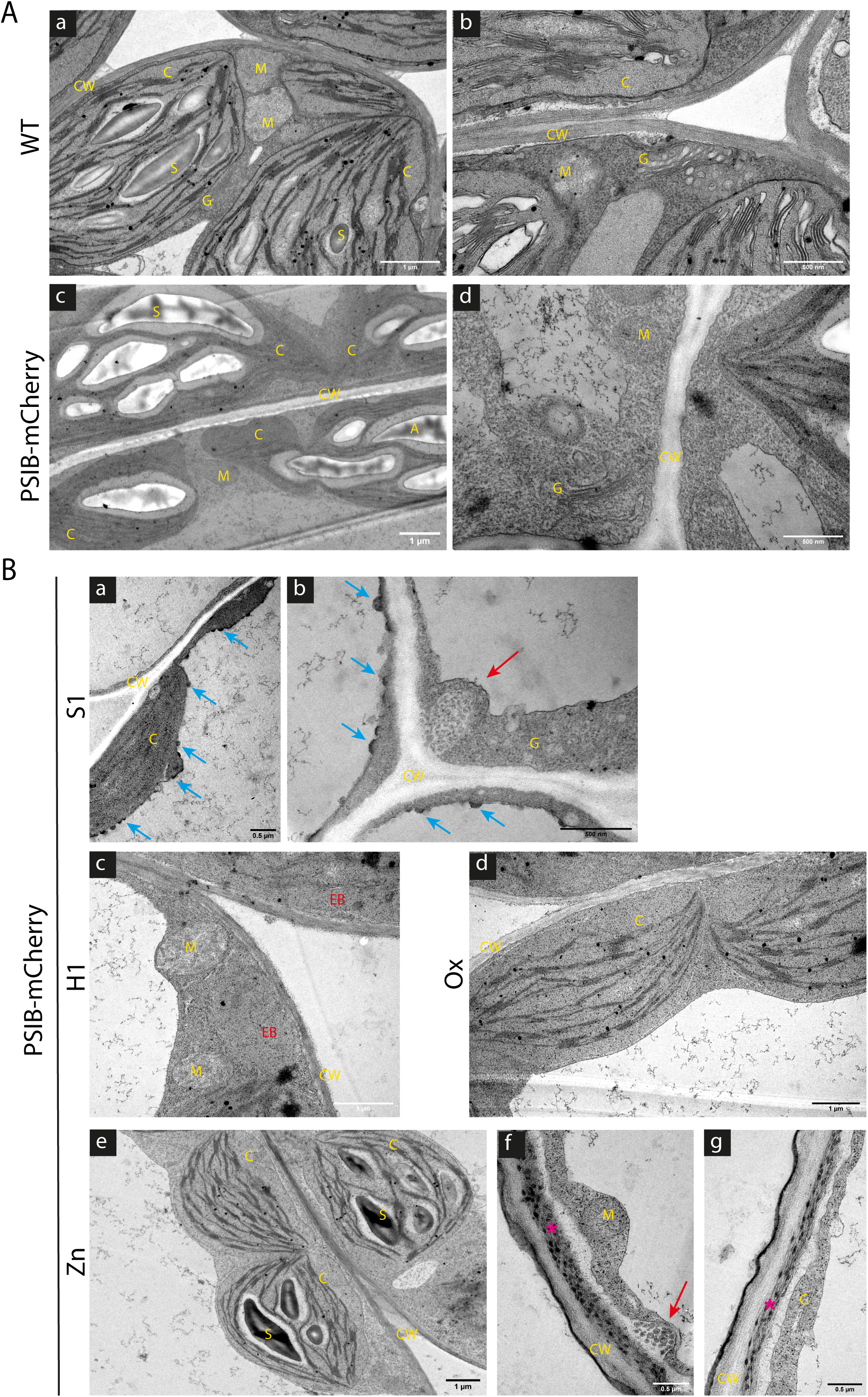
Micrographs revealing the ultrastructure of seedling cells from Arabidopsis thaliana wild-type and PSI B-mCherry line. **A** – Ultrastructural comparison between cells from WT and PSI B-mCherry line. The most relevant organelles are evidenced and labelled. Abbreviations: S, starch granules; C, chloroplasts; CW, cell wall; G, Golgi apparatus; M, mitochondria. **B** - Ultrastructure of cells from PSI B- mCherry line, under stress conditions. Saline stress (S1), mild hydric stress (H1), oxidative stress (Ox) and stress induced by metals (Zn). The most relevant organelles are evidenced and labelled. Red arrows indicate the vesicles and blue arrows indicate unknown electron-dense structures. Pink asterisks indicate possible zinc accumulations. Abbreviations: S, starch granules; C, chloroplasts; CW, cell wall; EB, ER body; G, Golgi apparatus; M, mitochondria. Scale bars: 0.5 µm. Images were analysed and processed using ImageJ/Fiji software.

Seedling cells from the PSI B line under saline stress conditions (S1) show significant compaction of the cytoplasm, which resulted in increased electron density, as shown in Figure 5-B (a,b). An intriguing feature observed for this condition was the presence of small electron-dense structures along the tonoplast, facing the lumen of the vacuole (Figure 5-B a,b; blue arrows). Moreover, some cells presented high number of vesicles clustered near the cell wall, possibly involved in secretion of material (Figure 5-B b, red arrow). Under hydric stress (H1), it was observable an increase in cytoplasm density when compared to the control condition (Figure 5-B c). It is important to highlight the lack of starch granules in the chloroplasts, a morphological alteration frequently associated with stress. The image also reveals mitochondria with typical ultrastructure and ER bodies (Figure 5-B c). Similarly to what was observed in saline (S1) and hydric stress (H1), in Ox condition there was evident alterations in the arrangement of thylakoids, specifically within the chloroplast lamellae, along with a substantial decrease in starch grain count (Figure 5-B d). Aside from that, the cytoplasm condensation was also noticeable along. Lastly, in the metal-induced stress condition (Zn), the chloroplast contain large starch grains, as observed for the control condition, when pointing our focus to the internal structure of the chloroplasts, it became apparent that the grana/intergrana complex was elongated (Figure 5-B e). Moreover, electron-dense structures can be found along the cell wall in some cells, hypothesized as corresponding to Zinc accumulations (Figure 5-B f, g; pink *). Apart from this condition, these structures were undetectable in any other situation. Also, on some occasions, clustering of vesicles near the cell wall were visible, similar to what was described for the S1 situations, possibly corresponding to secretion events (Figure 5-B f; red arrow).

### 3.5 PSI B trafficking pathways induced by stress

After analysing the control condition, it was previously shown (Figure 2-A) that, apart from its localization in the vacuole, PSI B-mCherry was identified in ER-derived elongated bodies in cotyledon cells (Figure 2-A, d), indicating the presence of PSI B in ER bodies. Concerning stress conditions, in S1, along with its prominent localization in the vacuole, PSI B-mCherry was once again detected in elongated structures (Figure 6-A, a-b, blue arrows). Under both hydric stress conditions (H1 and H2), there was notable vacuolar accumulation, along with a substantial presence of the elongated structures (Figure 6-A, c-f, blue arrows). PSI B-mCherry accumulation persisted in both the vacuole and elongated structures under the remaining conditions, oxidative (Ox) and metal-induced (Zn) stress (Figure 6-A, g-j, blue arrows). The number of these elongated structures seemed to be higher in the stress conditions when compared to the control, with the exception of the saline stress condition. Additionally, the movement of the ER-bodies in plants under stress was observed during time-lapse experiments (Figure 6-B). Interestingly, it was noticed that under hydric stress conditions (H1 and H2) the movement observed in control conditions was not noticeable (Figure 6-B, a-b, blue circles; Sup movie 4 and 5), while in Ox and Zn situations the movement of ER-bodies was still noticeable, despite apparently slower than control (Figure 6-B, c-d, blue circles; Sup movie 6 and 7).

**Figure 6 –.**
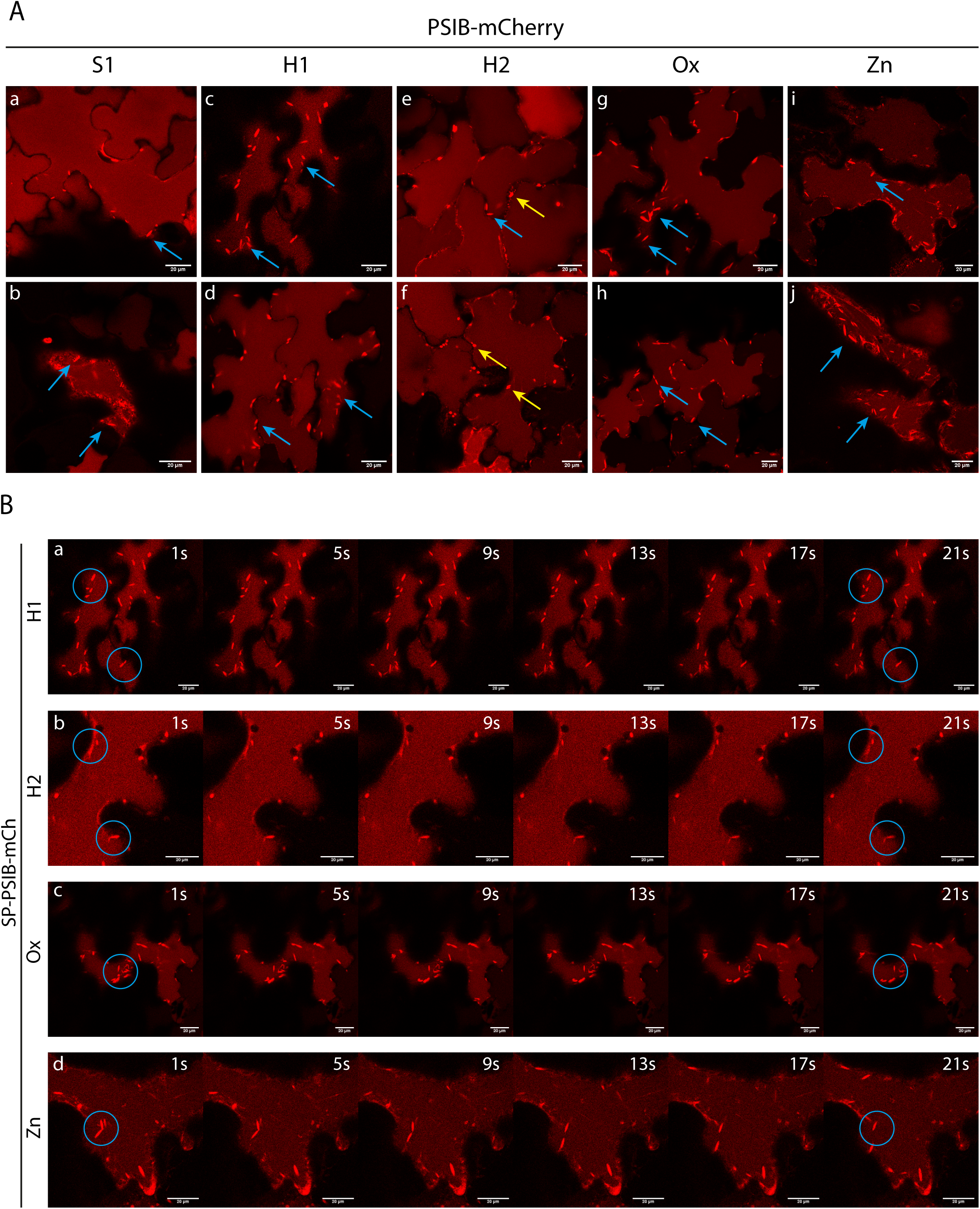
PSI B-mCherry localisation in Arabidopsis thaliana seedlings under stress. **A** - Expression of PSI B-mCherry in saline stress (S1), mild hydric stress (H1), high hydric stress (H2), oxidative stress (Ox) and stress induced by metals (Zn). Note the evident vacuolar accumulation in all tested conditions. The yellow arrows indicate small accumulations and blue arrows indicate ER bodies. Images were acquired using the 561 nm laser for mCherry and were analyzed and processed using ImageJ/Fiji software. **B** - Stills retrieved from time-lapse movie illustrating the movement of PSI B-mCherry-labelled compartment (blue circle) in mild hydric stress (H1), high hydric stress (H2), oxidative stress (Ox) and stress induced by metals (Zn).

### 3.6 Gene expression analysis

To evaluate the expression of PSI B and genes related to the vacuolar transport system, a qPCR analysis was performed (Figure 7). Even though the PSI B-mCherry transgene used in this study is driven by the 35S promoter, thus corresponding to an overexpressing line, the expression of the transgene was quite altered when the plants were under stress conditions. The results obtained for PSI B expression indicated an up-regulation in S1, Ox, and Zn conditions, while a down-regulation in H1 was observed, when compared to control situation (Figure 7-A). In the H2 condition, there is a slight upward trend, but with no significative changes. Intrigued by the analysis of PSI B expression results, we moved on to examine genes associated with the vacuolar transport system. The purpose was to determine if the expression of these chosen genes would be impacted in PSI B-transformed plants and could be related somehow with the changes observed for PSI B. We selected three Protein Storage Vacuole (PSV) trafficking-related genes (*AtRMR1*, *AtVTI12*, and *AtVSR1*), and two Lytic Vacuole (LV) sorting-related genes (*AtVSR2* and *AtVTI11*). Additionally, and due to the localization of PSI-B in ER-bodies, we also tested the expression of *AtVAMP723*, a gene coding for an ER-localised SNARE. The expression pattern of *AtVSR1* and *AtRMR1* is quite similar, with both being down-regulated in the S1 condition and up-regulated in Zn situation, while no changes were observed for the other vacuolar receptor tested – *AtVSR2* (Figure 7-B). Likewise, no significant changes were observed for *AtVTI12*, while *AtVTI11* is up-regulated in the Zn condition (Figure 7-C). For *AtVAMP723* the up-regulation in Zn condition is also clear, along with a down-regulation of this gene in the S1 condition (Figure 7-C). It is worth to note that up-regulation observed in the Zn condition for these genes is 4-6-fold higher than in control condition.

**Figure 7 –.**
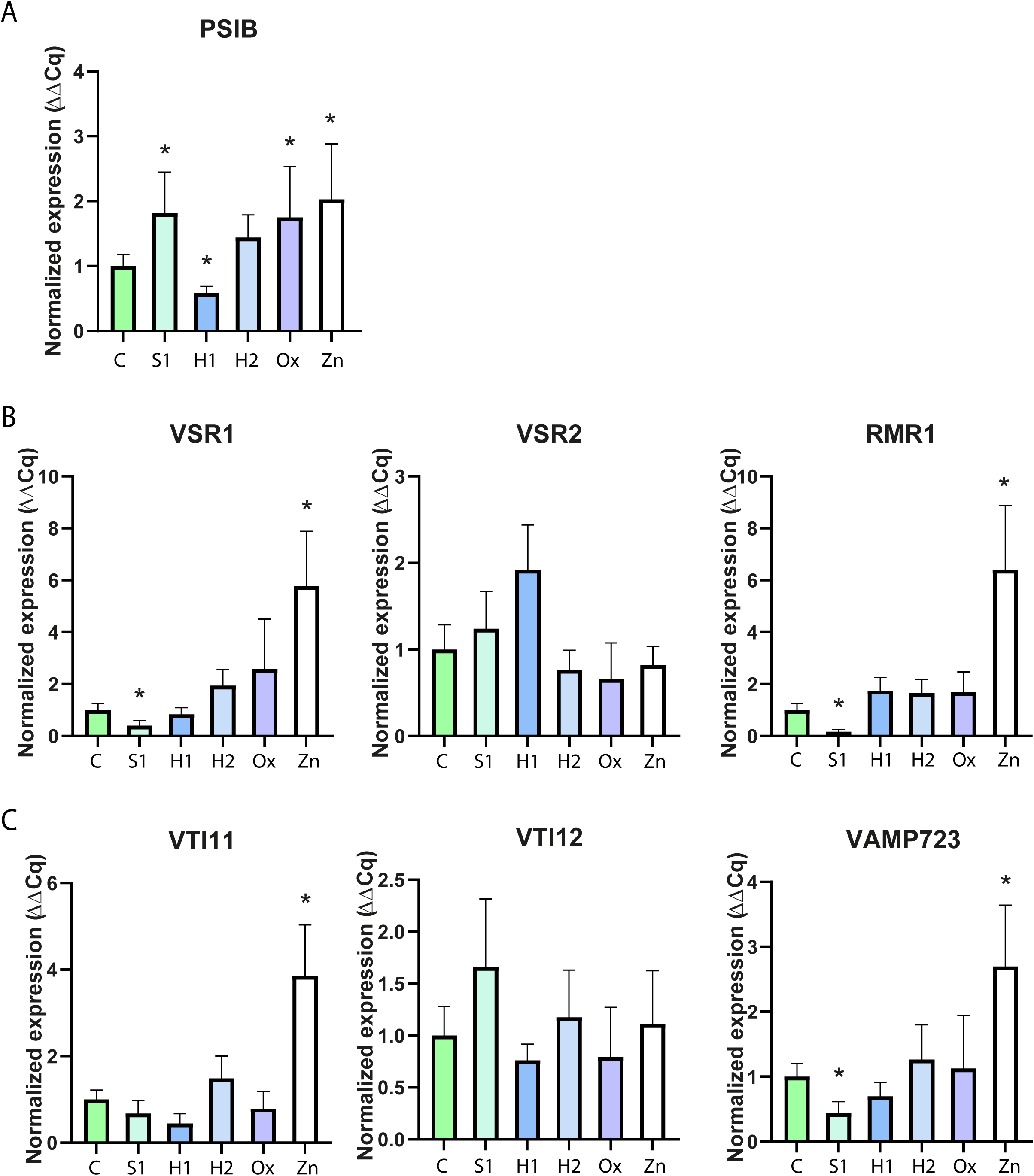
Expression by qRT-PCR in PSI B transformed plants under stress conditions. **A** - Bar chart analysis of PSI B gene in different stress conditions relative to control. **B** - Bar chart analysis of genes involved in the vacuolar transport system. Bar chart analysis of the vacuolar sorting receptors AtVSR1, AtVSR2, and AtRMR1 expression, in different stress conditions relative to control. **C** - Bar chart analysis of the SNAREs AtVTI11, AtVTI12, and AtVAMP723 genes, in different stress conditions relative to control. The statistical experimental from One-way ANOVA values are represented by: * (pvalue ≤0.05), according with the One-way ANOVA.

## 4. Discussion

Environmental responses are coordinated by many signaling pathways. When activated, these pathways trigger a chain reaction of events, including changes in transcription and translation, post-translational modifications, and regulation of protein degradation rate [35]. The communication process within the endomembrane system is essential for proper membrane trafficking and play a vital role in environmental responses. Cardosins A and B, aspartic proteinases present in thistle flowers, are recognized model systems for studying intracellular trafficking and are responsive to stress conditions [36]. In its turn, the PSI domain has frequently been associated with membrane reorganization and remodeling, and also vacuolar sorting [18,27,37]. It is also recognized that stress disrupts the conventional targeting of proteins to the vacuole, resulting in pathway shifts, sometimes bypassing the Golgi [14,38]. The major strategy here to investigate these pathway shifts was to obtain Arabidopsis plants overexpressing PSI B coupled to mCherry in order to understand the role of unconventional protein trafficking and how it might be affected under stress conditions.

### 4.1 New Insights into PSI B subcellular localization

To explore the subcellular localization of PSI B, we investigated cotyledons and leaves. In *Cynara cardunculus*, the native system, Cardosin B is detected in the extracellular matrix of the flowers’ stigma transmitting tissue [39], while in Arabidopsis and tobacco mature leaves (heterologous systems), it is accumulated in the LV [40]. The observation of PSI B accumulation in cotyledons primarily unveiled its presence in the vacuole. Nevertheless, it was also identified in the ER, likely representing an intermediate step in its route toward the vacuole. Additionally, the presence of PSI B within elongated aggregates is a notable and unprecedented observation. These structures, appearing to be ER bodies, have never been associated with PSI B accumulation before, nor with aspartic proteinases or other PSI domains. Moreover, the fact that ER bodies developed in the epidermal cells of the non-transgenic Arabidopsis cotyledons, reveals that those structures are not artificially caused by transgene overexpression [41]. It is crucial to emphasize here that ER bodies are structures directly derived from the ER that fuse with vacuoles without the involvement of the Golgi, and thus can be considered to be involved in an unconventional route [13,41]. Considering that PSI B was previously characterized as a mediator of the conventional route and being its transport dependent on COP-II vesicles [42], this represents a novel insight into our understanding of its intracellular trafficking, indicating a deviation from the established model for PSIs mediated trafficking. To provide a more comprehensive characterization of the PSI B’s localization, we explored its distribution in vegetative tissues, specifically in rosette leaves. It is noteworthy that while the majority of PSI B accumulation occurred within the vacuole, fluorescent signal was also detected in the ER, probably corresponding to molecules still in trafficking. The tissue-specific nature of PSIs (and aspartic proteinases in general) expression and trafficking has been long hypothesised as being associated with tissues under high metabolic activity, like flowers and seeds [37,43,44]. By contrast, in vegetative tissues low steady-state levels were observed in the native plant [45,46]. This specific localization of PSI B can thus be related with its role in seed tissues and is a good starting point for exploring its role in the plant, apart from its vacuolar sorting ability.

Along the different stress conditions under study, the ER bodies were still detected and apparently the number of ER bodies increased in stressed plants, correlating with our hypothesis that this particular localization must be related with a specific PSI function. Hayashi and colleagues (2001) documented that under saline stress conditions, seedlings exhibit a fusion of ER bodies with each other and with vacuoles, thereby mediating the direct delivery of proteinase precursors to the vacuole [13]. Another study found that wound stress and treatment with methyl jasmonate (MeJA) induced the formation of many ER bodies in rosette leaves, which had no ER bodies under normal conditions. As so, our results are consistent with prior research linking the formation of ER bodies to the response of the endomembrane system to environmental stress, thereby implying a biological role for ER bodies in the defense mechanisms of higher plants. ER bodies are distributed across the epidermis of cotyledons and hypocotyls in young seedlings but gradually disappear as the plant matures [41]. The findings regarding PSI B localization in leaves support this theory along with the idea that protein localization depends not only on the tissue in which it is expressed but also on the sorting pathways that can be triggered under stress [40,44,47].

Also remarkable was the results obtained after the use of drugs affecting the actin and microtubules cytoskeleton. The movement of ER bodies in plant cells is indeed linked to the actin cytoskeleton. Several studies highlight the interaction between the ER and actin in plant cells, and actin-membrane adaptors, such as NET proteins, have been identified as key players in mediating the interaction between the actin cytoskeleton and the ER membrane, facilitating the movement and anchoring of ER bodies within the cell [48,49]. Furthermore, specific proteins, such as formins, which regulate actin dynamics by functioning at the barbed end of actin filaments, have been found to localize to the nuclear envelope and interact with the ER, suggesting a complex interplay between these cytoskeletal elements and the ER network [48,50]. Our results fully support these past observations, but also show a similar phenotype in cotyledons under hydric stress. Very few studies address the role of the plant cytoskeleton under stress and how it can be affected, which are summarized in a recent review [38]. However, given the importance of actin filaments and microtubules in protein transport, cellular organization and organelle positioning, it is of greatest importance to study these effects. The results presented here do not allow to discern if the stress applied affect the actin itself or actin-associated proteins or motors, yet is a good starting point for further studies.

### 4.2 PSI B as a potential player in abiotic stress tolerance

Given the recent implication of aspartic proteinases and PSIs themselves in plants’ response to stress [51–53], we took advantage of the Arabidopsis line expressing PSI B to evaluate the extent of this hypothesis. The plant root system is critical in the plant’s response to various types of abiotic stress. The root system analysis of PSI B plants indicates a downward trend in root length for all conditions except H1. This behaviour aligns with a well-established pattern, as plants generally extend their root systems in response to osmotic or drought conditions to enhance water uptake [54–56]. The PSI B line shows a notable increase in average root length compared to the WT plants, evident in both control and H1 conditions. These findings are fascinating as they show that PSI B overexpression line revealed behaviour similar to non-stressed plants. In 2015, Guo and colleagues constitutively expressed a grape aspartic protease gene (*AP17*) in transgenic Arabidopsis, revealing significantly bigger root systems compared to the WT [57]. A similar outcome was reported when *APA1* was overexpressed in Arabidopsis, leading to an increase in root length in the transformed plants [58]. As a result, our data on PSI B correlates with previously findings in the literature, indicating that this domain may play a role in enhancing plant tolerance to mitigate the impacts of stress conditions, particularly osmotic stress. On the other hand, throughout the stress trials, it was clear that the seedlings’ leaf area was substantially compromised. Drought and salinity are both known to have negative impacts on crops, most notably inducing a decrease in leaf area [59] corresponding to the conditions where we observed the major alterations. Given these results, particularly in what concerns plant development and leaf area and because abiotic stress have a major impact on the photosynthetic machinery [60], we decided to determine the levels of total chlorophyll and carotenoids. Regarding total chlorophylls, despite no significative changes observed, PSI B line show a slight upward tendency observed in all conditions except for Zn, behaving nearly like non-stressed plants. Carotenoids serve as accessory pigments synthesized by plants to assist chlorophyll in capturing light [61]. Observing the carotenoid levels in WT and PSI B plants, it was clear that the pattern differs from the one observed for chlorophylls. In this case, there was a noticeable upward trend in nearly all conditions, particularly in H1 and H2. Initially, these findings may appear contradictory, as we might anticipate a decrease in carotenoid content. However, it is important to note that carotenoids serve a key function in protecting plants from oxidative damage and are used by the plant’s defense mechanisms [62,63]. Consequently, their production tends to increase when the production of chlorophyll is compromised [32]. The chlorophyll results are thus related to the plant’s developmental response. The decline in chlorophyll levels induces a decrease in photosynthetic rate, resulting in a delay in plant development, which is particularly noticeable under osmotic stress conditions. In metal-induced stress, the decrease in chlorophyll is primarily manifested as chlorosis in the leaves. This enables the plant to reallocate resources to other areas, such as the roots, resulting in a reduction in the photosynthetic rate without necessarily compromising overall growth. Which also explains why root length is not affected as much compared to other conditions.

To gain a more comprehensive understanding of possible modifications induced by the overexpression of PSI B, the ultrastructure of Arabidopsis seedlings was assessed using TEM. Compared to WT, PSI B transformed plants in control condition, maintained chloroplast structure and organization. The ultrastructure of WT cells under the stress conditions studied has already been characterized in a recent study [33]. The first aspect that caught our attention regarding the PSI B line was the compaction of the cytoplasm in S1 condition, which was not observed in WT plants. Also, thylakoid dilatation was visible as well as a decrease in both the number and the size of starch grains, resulting in a change in the chloroplast structure. This decrease is attributed to the hydrolysis of starch granules, which serve as a source of monosaccharides that help mitigate the effects of salt stress. Both of these responses occurred in WT and correlate with previous studies [33]. The most striking difference from the WT cells under the same conditions, and from the other conditions observed are the small electron-dense structures decorating the tonoplast, that remains to be determined. In the H1 condition, the chloroplasts were devoid of starch grains, which differ substantially from those reported by Neves *et al*. (2021), who identified large starch granules [33]. In Ox condition changes in chloroplasts’ organization were observed, particularly in the arrangement of the thylakoids, which are likely the result of lipid peroxidation, a process that affects the membrane’s integrity [64]. Given the essential role of thylakoids in photosynthesis, a decrease in photosynthesis activity can be predicted. Upon exposure to metal-induced stress, a unique occurrence was the presence of electron-dense structures within the cell wall. In contrast, when WT plants were exposed to the same condition, there is no evidence of these structures [33]. The discovery of such structures, together with the analysis of PSI B expression may suggest an involvement of PSI B in the accumulation (or translocation) of Zn in the cell wall and vacuole as a detoxification mechanism. The PSI B overexpression does not promote major alterations in the cell architecture, when comparing to WT plants under the same conditions, yet there are some specific features that indicate some difference and may be related to a PSI B putative role in plants under stress.

### 4.3. From stress signals to gene expression: the enigma of plant response

To have further insights on the effect of stress signals on Arabidopsis plants overexpressing PSI B, we examined the expression levels of PSI B, and genes related to the vacuolar transport system. Despite being driven by 35S promoter, and based on the results obtained so far, we investigated whether there was some additional regulation of PSI B expression under stress conditions. In fact, there is some. PSI B was shown to be upregulated in S1, Ox, and Zn but not in hydric stress (H1 and H2). The upregulation of PSI B in the S1 condition may be associated with its function in membrane interactions [18,26]. This is particularly relevant considering that, in response to salt stress PSI B could be involved in helping the plant maintain proper ion balance through ion transport and osmotic adjustment. However, it is intriguing that the same was not observed in the hydric stress conditions, as the cellular response is similar. Oxidative stress leads to modifications to the redox status of the cells and it is a consequence of any stress imposed to plants [65]. Thus, being difficult to clarify the PSI B upregulation in this situation. However, the observed upregulation of PSI B under metal-induced stress presents a highly interesting result. It is well established that vacuoles serve as the primary sites for metal sequestration in plants [66–68]. Given the nature of PSI B as vacuolar sorting signal, this expression trend suggests that PSI B might play a role in the detoxification process of Zn by mediating its transport to the vacuole. This could also be related with PSI ability to interact with membranes and to promote vesicle fusion [18,69]. In any case it is a quite important finding to pursue and to uncover whether PSIs may have a role in metal detoxification.

To check whether alterations in the PSI B expression and localization were accompanied by alterations in the expression of genes involved in intracellular processes, we evaluated the expression of genes particularly related with vacuolar routes - *AtVTI12*, *AtRMR1*, *AtVSR1*, *AtVSR2*, and *AtVTI11*, and the ER-associated *AtVAMP723*. A recent study conducted in our laboratory showed that WT plants under abiotic stress conditions presented differential gene expression, suggesting that stress poses a positive regulatory role in the trafficking routes [33]. *AtVSR2* is a gene coding for a vacuolar sorting receptor involved in clathrin-coated vesicles sorting from the Golgi apparatus to the LV [70] and the results showed that under stress conditions, its expression in PSI B plants remained reasonably stable. This effect was not observed in WT plants, under the same conditions, as this gene was downregulated in all stress conditions [33]. On the other hand, *AtVSR1* and *AtRMR1* showed a similar response, being under-expressed in salt stress and overexpressed in Zn condition. This pattern in S1 condition contrasts with what was observed in WT plants, where these genes were overexpressed under osmotic and metal induced-stress [33]. In fact, *AtVSR1* (like *AtRMR1*) codes for a vacuolar-sorting receptor involved in clathrin-coated vesicles sorting from the Golgi apparatus to vacuoles [71] with an established role in the osmotic stress response [72]. The change in its expression in plants overexpression PSI B is another hint on the putative defensive role of PSI B implying that it may function as a positive regulator or enhancer of stress-responsive genes.

Regarding the SNAREs coding genes analyzed during this assay, no major changes in expression were observed in PSI B seedlings under stress, with the exception for the overexpression of *AtVTI11* and *AtVAMP723* in Zn condition. This increase in expression in the metal-induced stress is quite consistent between the genes tested – PSI B, *AtRMR1*, *AtVSR1*, *AtVTI11* and *AtVAMP723* – and cannot be ignored, as it suggests that PSI B might play a role in somehow promoting gene expression under specific stress conditions or may have a protective role or an enhancing effect on gene expression under stress. Therefore, we consider that PSI B may likewise play a role in the cellular response to metal detoxification. This could involve modifying membrane permeability and facilitating metal accumulation in the vacuole, which could be possible given the membrane-binding and modulation abilities of this domain [69,73]. Also intriguing is the behavior of *AtVTI12*, which expression does not differ from control in PSI B plants, but is extremely overexpressed (10-30-fold higher than control) in WT plants under the same conditions [33]. AtVTI12 belongs to a class of SNARE proteins and is involved in the trafficking of proteins to the PSV [74,75], being also involved in the docking process of autophagic and exocytic vesicles with the vacuole [76]. Despite being clear an impact of PSI B overexpression in the expression of *AtVTI12* gene, even if indirect, at this point is not possible to ascertain which pathway PSI B is affecting. Considering all the data, numerous assumptions and unanswered questions remain. It is clear that PSI B overexpressing can, on its own, impact plant sorting pathways, resulting in changes in gene expression. Moreover, given PSI B’s involvement in the plant’s stress response, its overexpression may disrupt the protein sorting system and influence the expression of other stress-responsive genes.

## 5. Conclusion

In response to abiotic stress conditions, certain physiological and cellular changes take place within plant cells. Plants have evolved specific adaptations involving adjustments in protein production, trafficking, and endomembrane remodeling to cope with these challenging conditions. Therefore, understanding trafficking mechanisms as well as the proteins involved becomes essential for uncovering the plant’s response to stress. This study successfully led to the establishment of stable Arabidopsis lines overexpressing PSI B, providing a valuable resource for the laboratory’s future projects that have been investigating the APs trafficking pathways and the mechanisms mediated by these Vacuolar Sorting Determinants (VSDs). The findings presented suggest that PSI B play a significant and active role in enhancing plant fitness, revealing its value in adaptation and tolerance to abiotic stress. Moreover, the identification of PSI B accumulation in ER bodies is an unquestionably novel and significant discovery, along with the expression results observed in Zn-mediated stress condition, sparking a lot of new hypotheses. One interesting question that remains is if other PSIs could have the same stress-protective effect. It has been documented that PSI can enhance the tolerance of plants to the fungi *Botrytis cinerea* [23] and that it has antimicrobial activity on plant and human pathogens [20]. More recently, a review from Cheung and collaborators [51] explores the relevance of PSIs in defense mechanisms, highlighting its structure, conservation among species, mode of action and diverse trafficking routes, as key features to fully understand their functions and potential applications in enhancing plant resistance to pathogens. Other question that remains unclear, and this one being more difficult to address, is the fact that PSI domain is synthetized as part of a proteinase, and is removed along the proteinase processing. Does the domain, after cleavage, also presents the same properties as when expressed isolated? While much remains to be explored, the findings presented here represent a promising start towards deciphering the mysteries of changes in protein trafficking induced by stress.

## 6. Supplementary data

<u>Supplementary Figure 1</u> - Subcellular localization of PSI A-mCherry in *A. thaliana* overexpression plants. No ER-bodies are detected in cotyledons of Arabidopsis seedlings expressing PSI A, contrary to what is shown for PSI B Arabidopsis line.

<u>Supplementary Figure 2</u> - Direct comparison of biometric parameters and pigments of both *Arabidopsis thaliana* WT and of PSI B-mCherry line, under abiotic stress.

<u>Supplementary Movie 1 –</u> Time-lapse experiment showing the PSI-B labelled ER-bodies moved along the ER network.

<u>Supplementary Movie 2 –</u> Time-lapse experiment showing the PSI-B labelled ER-bodies moved along the ER network after Cytochalasin D treatment.

<u>Supplementary Movie 3 –</u> Time-lapse experiment showing the PSI-B labelled ER-bodies moved along the ER network after Oryzalin treatment.

<u>Supplementary Movie 4 –</u> Time-lapse experiment showing the PSI-B labelled ER-bodies moved along the ER network in plants under hydric stress H1.

<u>Supplementary Movie 5 –</u> Time-lapse experiment showing the PSI-B labelled ER-bodies moved along the ER network in plants under hydric stress H2.

<u>Supplementary Movie 6 –</u> Time-lapse experiment showing the PSI-B labelled ER-bodies moved along the ER network in plants under oxidative stress (Ox).

<u>Supplementary Movie 7 –</u> Time-lapse experiment showing the PSI-B labelled ER-bodies moved along the ER network in plants under Zn conditions.

## 7. Author contributions

Conceptualization – AS, CP; Formal analysis – IM, JN, CP; Funding acquisition – JP, CP; Investigation – IM, JN, CP; Methodology – IM, CP; Project administration - CP, JP; Resources – JP, CP; Software – IM, CP, JN; Supervision – CP, AS, SP; Validation – CP, JP, AS, SP; Roles/Writing - original draft – IM, CP; and Writing - review & editing – IM, JN, CP, JP, AS, SP.

## 8. Conflict of interest

The authors declare no conflict of interests.

## 9. Funding

Research funded by national funds via FCT (Foundation for Science and Technology) through the Strategic Projects UIDB/05748/2020 and UIDP/05748/2020, DOI https://doi.org/10.54499/UIDP/05748/2020 and https://doi.org/10.54499/UIDB/05748/2020.

## 10. Data availability

All data is available upon request.

## Notes

### Competing Interest Statement

The authors have declared no competing interest.

